# How many samples are needed to infer truly clonal mutations from heterogenous tumours?

**DOI:** 10.1101/606053

**Authors:** Luka Opasic, Da Zhou, Benjamin Werner, David Dingli, Arne Traulsen

**Author notes:** Corresponding author: Arne Traulsen.

## Abstract

**Background:** Modern cancer treatment strategies aim to target tumour specific genetic (or epigenetic) alterations. Treatment response improves if these alterations are clonal, i.e. present in all cancer cells within tumours. However, the identification of truly clonal alterations is impaired by the tremendous intra-tumour genetic heterogeneity and unavoidable sampling biases.

**Methods:** Here, we investigate the underlying causes of these spatial sampling biases and how the distribution and sizes of biopsies in sampling protocols can be optimized to minimize such biases.

**Results:** We find that in the ideal case, less than a handful of samples can be enough to infer truly clonal mutations. The frequency of the largest sub-clone at diagnosis is the main factor determining the accuracy of truncal mutation estimation in structured tumours. If the first sub-clone is dominating the tumour, higher spatial dispersion of samples and larger sample size can increase the accuracy of the estimation. In such an improved sampling scheme, fewer samples will enable the detection of truly clonal alterations with the same probability.

**Conclusions:** Taking spatial tumour structure into account will decrease the probability to misclassify a sub-clonal mutation as clonal and promises better informed treatment decisions.

## Background

In the past years, it has become increasingly clear that cancers are typically highly heterogeneous and characterised by a large degree of spatial diversity, which complicates cancer therapy [1, 2]. Modern anticancer therapies aim at targeting tumour-specific genetic and epigenetic alterations, e.g. by specifically designed molecules [3] or immuno-therapy [4–6]. The paradigmatic example has been Chronic Myeloid Leukemia that is driven by the BCR-ABL oncogene. Tyrosine kinase inhibitors (TKI) such as Imatinib can inhibit the critical gene driving the disease leading to long lasting remissions, improved survival and perhaps even cure in some patients [7–10]. With rare exceptions [11], this goal to date has not materialized for other tumours since in many of them, the appropriate driver mutations(s) are either unknown, not targetable or treatment resistant. Although now tumour sequencing is available commercially and mutations within tumours can be identified routinely, the clinical benefit of such therapies has been limited, since it is likely that the identified mutations are not responsible for driving the tumour in that specific patient [12], or resistance emerges fast after an initial brief treatment response [13–17]. It is important to note that simply because a mutation is ‘common’ in a specific tumour type does not make it an appropriate target of therapy. Determining which mutations are targetable is not simple for a variety of reasons including (i) the identified mutations may not be drivers in that patient, (ii) more than one driver mutation may be present, (iii) genetic and spatial heterogeneity within the tumour make it difficult to be reasonably certain that the truncal/clonal mutations that could be targeted have been properly identified, which is the main focus of our work here.

The accumulation of mutations in a growing tumour leads to the presence of cells with different mutational profiles. Classical branching models predict that mutations will be increasingly present at lower frequency [18]. More specifically, late alterations are typically found in a small proportions of cells, whereas early alterations are expected to be more abundant [19]. For example, mutations present in the first tumour initiating cancer cell should, in principle, be clonal and consequently found in every cancer cell of the particular patient. In clinical protocols, it is often assumed that a mutation that is present in approximately 50% of the sequencing reads of a single tumour bulk sample is likely clonal (after adjusting for tumour ploidy and tumour purity). However, this reasoning is problematic, since it requires that the underlying sample is representative of the whole tumour. Multiregion profiling shows that this is not the case [1, 2, 20]. These multi-region sequencing studies have revealed a much more complicated picture of severe inter- and intra-tumour genetic heterogeneity [1, 2, 21–23]. Mutations that appear clonal in a single sample can be sub-clonal or even absent in other samples of the same tumour [24]. Therefore, targeting such mutations would not be expected to provide long term therapeutic benefit as we would at best treat only the part of the tumour that contains these sub-clonal mutations.

Indeed, determining which are the truly clonal alterations in a neoplasm that contains billions of cells distributed in complex spatial patterns is a challenging problem that has important implications for modern cancer therapies. Limitations of sequencing depth, genetic heterogeneity within single samples, contamination with healthy tissue, and loss of genetic elements due to genome instability all complicate the classification of these alterations [25]. In this regard, multi-region sequencing has been shown to be more informative for the discrimination of mutations than single bulk sequencing [1, 18, 26–29]. As multi-region sequencing of tumours has become feasible, this has led to the development of a range of phylogenetic methods and tools to construct phylogenetic trees from cancer and infer truncal (clonal) mutations [30, 31].

The detailed architecture of any tumour is likely unique and driven by the complex interactions between microevolution, the immune response as well as the presence of physical barriers to growth of the tumour population in each specific patient. The latter depend on the location of the tumour within the body. This complexity makes it difficult to reconstruct the branching process that underlies the growth of the tumour population. In the absence of such knowledge, what would be the optimal sampling approach for each individual tumour and how can we maximize our probability to identify truly clonal (and hopefully driver) mutations within these tumours. This is the focus of our work here.

We have previously shown that it is not necessary to reconstruct the complete phylogenetic tree of a tumour to estimate the probability to identify all clonal alterations correctly [20]. It is easier and sufficient to identify only the earliest branching events, which then allows the detection of all truly clonal genetic alterations within individual tumours. We consider the earliest branching event that separates sub-clonal mutations from those mutations present in the ancestral population of cancer cells. We refer to these branch-defining sub-clonal mutations on the ancestral population background as first-tier mutations (Figure 1(d)). Taking more samples will naturally increase the chance to exclude misclassified sub-clonal mutations. However, this obviously implies a cost-benefit tradeoff and a proper understanding of the scaling of these probabilities with increasing tumour sample numbers can better inform treatment strategies.

In our previous work, we showed how the probability to correctly classify clonal mutations scales with additional samples and how many samples are needed to identify the truncal mutations with a certain level of confidence given the life histories of a tumour [20]. But critically, this initial study did not consider successive branchings from the ancestral population, the spatial structures of both the tumour and the samples as well as the influence of the sizes of the biopsies taken.

Here, we quantify the probability to correctly classify the clonal mutations of individual tumours growing in space with sufficiently low mutation rates per cell division (as is for example the case for clinical targeted or exome sequencing protocols). We also show how the size of tumour samples influences our ability to identify truly clonal alterations and how we can increase the accuracy of the detection by exclusion of low frequency mutations. We address the problem of spatially structured tumours, which can have great repercussions on clonality inferences. Finally, we compare different sampling protocols by comparing standardized spatial sampling patterns against random sampling. By applying standardized sampling patterns one can further increase the probability to correctly classify truly clonal mutations.

## Methods

### Computational model of tumour heterogeneity

We simulate tumours on a lattice where filled nodes represent the presence of individual tumour cells. For the structured case, the neighbouring cells are thought to represent a real – but arguably highly idealized – tumour architecture. For the unstructured case – often refereed to as the “well mixed” case – the neighborhood on the lattice has no relevance for simulations, as any cell can divide and place its offspring to any empty site. Simulations start by transforming one cell into a tumour cell through the introduction of the first mutation (in principal many mutations could be introduced, however this does not matter for our purpose here). Simulations run in discrete time steps. During each time step, one cell is chosen randomly for reproduction. Once chosen, it divides into one adjacent empty space, if the tumour is spatially structured, or in any random available place chosen from non-occupied spaces, for a well-mixed tumour. With each division, one daughter cell accumulates a novel mutation proportional with probability *µ*. After division, one randomly chosen tumour cell will die with a probability *d* = 0.1. To simulate a non-aggressive tumour, we have chosen a lower death rate than suggested by Waclaw et al. [32] (*d* = 0.5*b*) for the simulation of highly aggressive tumours.

Computationally, the tumour is represented by a sparse matrix, wherein the position of a cell, the ID of its parent cell and the signature identifier of each new mutation is stored. This information allows us to reconstruct the mutational profiles of any cell at any given time point. We assume that each mutation can arise only once during division and can only be lost when the a cell dies (corresponding to the infinite allele assumption). Moreover, we assume all mutations to be neutral – they do not affect the fitness of the carrier cell. Our assumption of neutrality should not impact the generalizability of our results. After a full sub-clonal sweep, the dominant sub-clone would appear as ancestral population, thus thus leading to a tumour population with similar underlying branching structure. The nature of our simulation makes the structured tumour grow mostly at its periphery. Once the centre of the tumour is densely populated cells can only divide if neighbouring space becomes available after a random cell death. This pattern is supported by observations of similar peripheral growth patterns in some real tumours [33].

In our analysis only the presence of new mutations is important and not the number of new mutations in each cell. We therefore assume that during each division daughter cells receive a new mutation with probability *µ* = 0.5. This assumption is supported by the estimated high mutation rates in neoplasms. With effective mutation rates (mutation per surviving lineage) of up to 10^*−*7^ mutations per base per division, we can expect a mutation occurring during almost every division within the exome of cancer cell [19]. With this mutation rate we achieve early branching and extensive intra-tumour heterogeneity from the early stages of tumour growth. This feature of the model is compatible with the fact that only early mutations are likely to spread sufficiently to be detected by Next Generation Sequencing [29]. Early branching provides a broad spectrum of sizes of the first sub-clone in our simulations due to the stochastic nature of sub-clonal growth and mutation accumulation. At the end of the simulation we calculate the frequency of each mutation within the tumour. We specifically denote frequencies of first tier sub-clonal mutations and the most frequent sub-clonal mutation. In the evolutionary history of the tumour, we define the first-tier branch as the subpopulation of cells that diverged directly from the ancestral population of tumour cells. To reconstruct the truncal mutations from multi-region tumour sampling, we need to either sample from two different first-tier subpopulations that emerged from the ancestral cancer population or from one first-tier subpopulation and the ancestral subpopulation, because sampling from the same first-tier branch will falsely identify branch-defining mutations as clonal mutations.

### Sampling and clonality inference

A single simulated biopsy is composed of a group of cells in close proximity (Figure 4), or a single cell (Figure 2) initially taken from a random location of the lattice. In a well-mixed tumour, due to the absence of spatial structure, a sample is a number of randomly pooled cells (Figure 4), or a single cell (Figure 2). We reconstruct the mutational profiles of the sampled cells and calculate the frequencies of the mutations within each sample. As we are unable to detect low frequency mutations with current sequencing technology (sequencing depth threshold), we vary the threshold *ε* to detect a mutation within the sample. Mutations that appear clonal across a tumour are those mutations present in all taken samples. However, in our simulations we know the ground truth and we can test how often these mutations actually represent truly clonal mutations present in the first cancer initiating cell. If no mutations were wrongly classified as clonal we mark our sampling as correct. Otherwise, if there is at least one sub-clonal mutation misclassified as clonal, we consider our sampling incorrect. To get the proportion of correct estimations for single tumours, we repeat the sampling process 10 000 times with *n* samples (shown as dots in Figures 2, 4).

**Figure 1:**
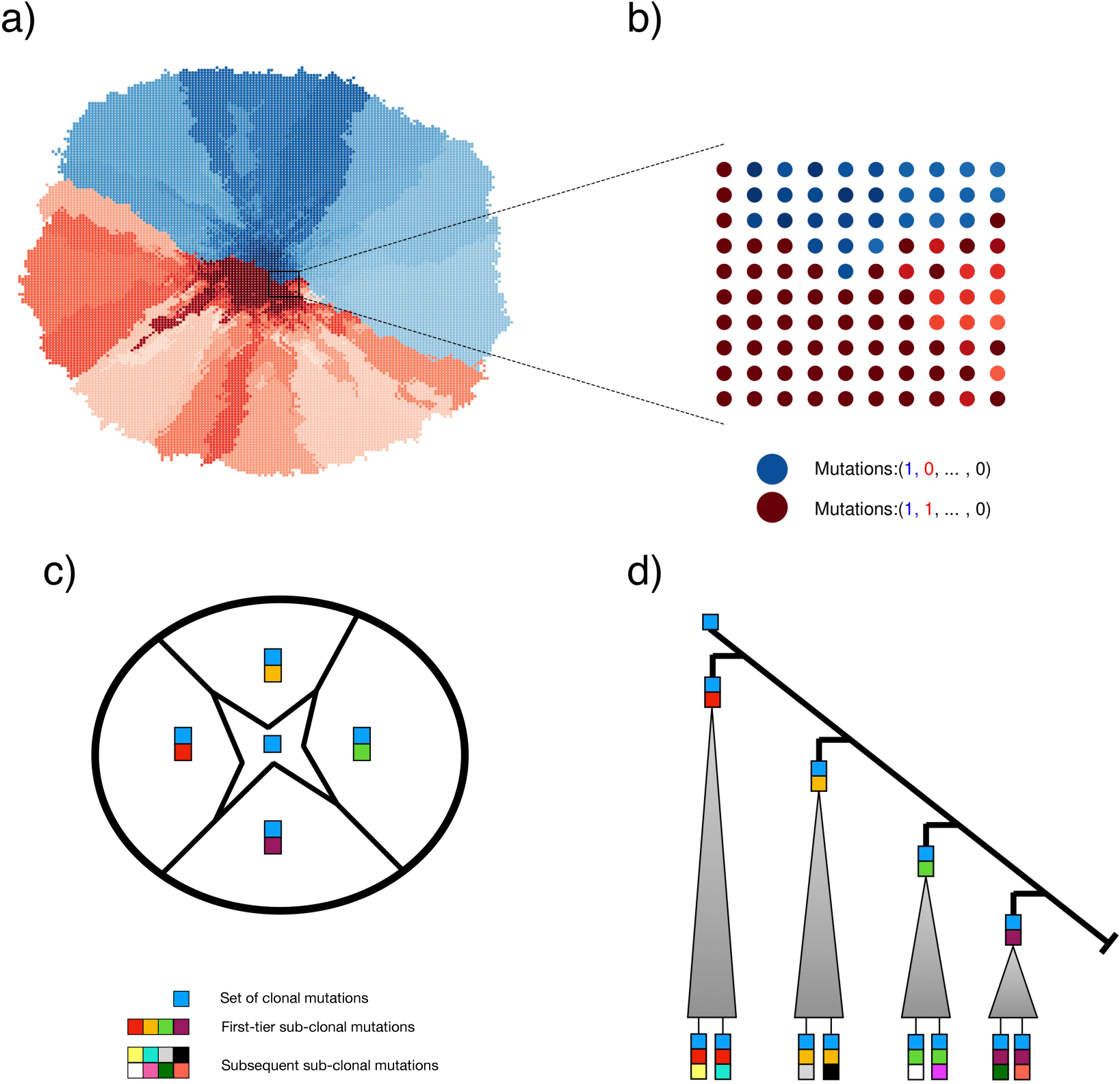
Spatial model of intratumour heterogeneity. **a)** Schematic illustration of our spatial cancer model. Tumour cells are represented as nodes on a two dimensional lattice. Each cell has a mutational profile. With each cell division, a cell can mutate with probability *µ* = 0.5 and thus intratumour heterogeneity is generated. While we typically think of each node as a cell, it could in principle also represent a small subpopulation of cells. **b)** The mutational profiles of cells within a bulk biopsy are then combined to give the mutational profile of the biopsy sample. Red colored subpopulations are cells that carry the first sub-clonal mutation (red in the mutational profile), while blue cells do not contain it. All cells carry one truncal mutation (blue in the mutational profile). Cells acquire new mutations and are presented with a different nuances of the original colour. In this exam-ple, because the red and blue subpopula4tions are approximately equal in size, we call the tumour well-balanced. **c)** The different first-tier sub-clones are spatially represented within the tumour, placed around the centrally positioned ancestral population. **d)** An ancestral population that contains a set of truncal mutations (blue square) branches multiple times into first-tier sub-clones with their own private sets of mutations. Having only samples from one of the major branches will result in a misclassification of the sub-clonal mutation that founded that branch.

**Figure 2:**
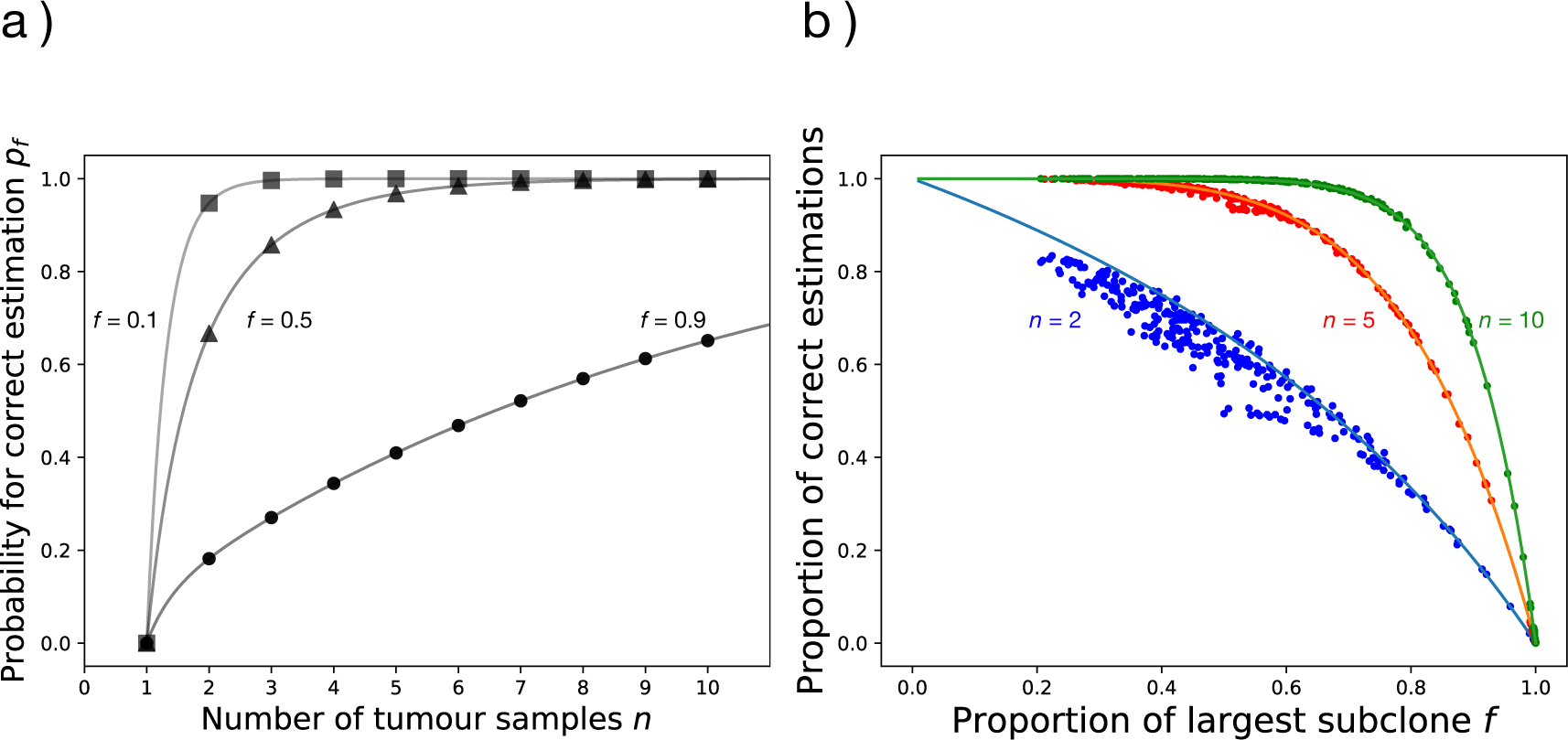
Comparison of clonality inferences in structured and unstructured models of tumours with a different proportion of the largest sub-clone. **a)**The probability to correctly identify set of truly clonal mutations with *n* tumour samples in our model. In tumours where the size of the largest sub-clone *f* is small (*f* = 0.1) the probability to correctly identify truly clonal mutations is already sufficiently (*>* 98%) high with two samples. In balanced tumours with *f* = 0.5, five samples give the same probability. **b)** The quality of our clonality estimation in dependence of the proportion of the first sub-clone. Lines represent solutions of the mathematical model, while dots represent results from clonality inferences of spatial computer simulations. A number of randomly distributed single-cell samples *n* (*n* = 2 shown in blue, *n* = 5 shown in red and *n* = 10 shown in green) was taken from each simulated tumour and clonality of present mutations was estimated. Each single dot represents the proportion of correct estimations for one tumour by sampling *n* tumour samples after 10 000 repetitions. Results from simulations are in agreement with model predictions for the full range of *f*.

**Figure 3:**
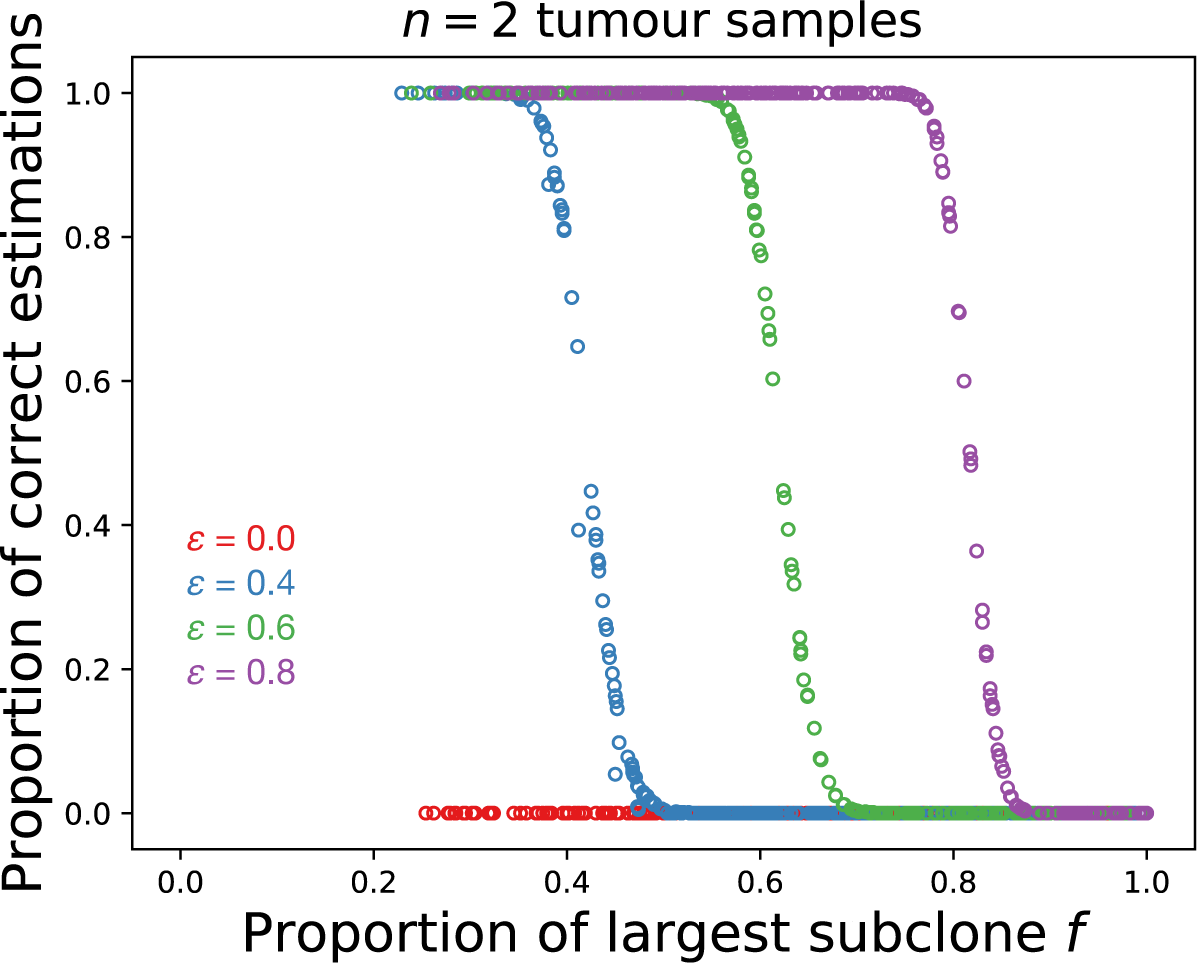
Effect of sample size on clonality inference in well-mixed tumour. The proportion of correct clonality estimates for *n* = 2 samples, each containing 1% of total number of cancer cells. *ε* represents the frequency bellow which we discard mutations from the clonality analysis. The value of *ε* where identification of clonal mutations becomes impossible corresponds to the proportion of the largest sub-clone *f*, as each sample is representative of the whole tumour.

**Figure 4:**
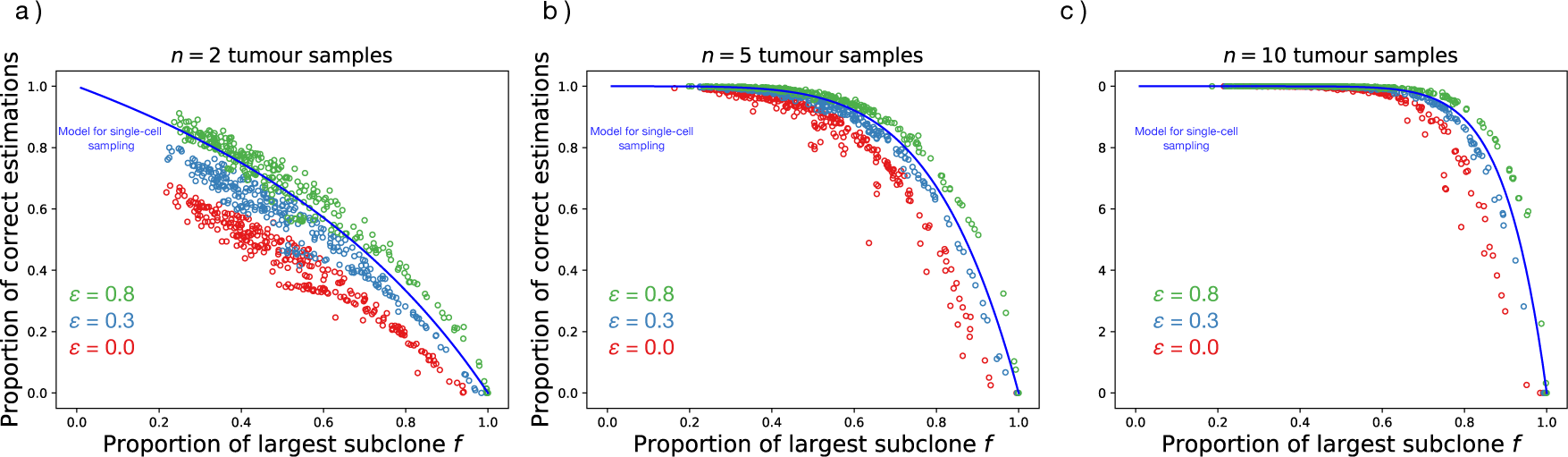
Effect of sample size on clonality inference in spatial tumour. To test the effect of biopsy size, we tested the accuracy of clonality estimations by sampling groups of cells (one sample = 200 cells, corresponding to 1% of tumour size). Batches of tumour samples (two for a), five for b) and ten for c)) were taken at random locations and used to estimate the clonality of the present mutations. We repeated sampling process 10 000 times for on each tumour and calculated proportion of correct estimations – single point on figure. We exclude mutations below a certain frequency *ε* from the analysis (*ε* = 0.0 or no exclusion for red, *ε* = 0.3 for blue, *ε* = 0.8 for green), which increases the accuracy of clonality estimations.

For pattern sampling we chose four single-cell samples from the tumour edge located at opposite directions and assessed clonality as previously described. To calculate the proportion of correct estimations for each individual tumour using our sampling pattern, we rotated the samples stepwise and assessed clonality on each step until we covered the whole tumour circumference (Figure 6(a)). Rotations of the samples allowed us to make multiple repetitions of sampling on a single tumour using the same pattern. The proportion of correct estimations was then compared with random single-cell sampling and our mathematical model quantification.

**Figure 5:**
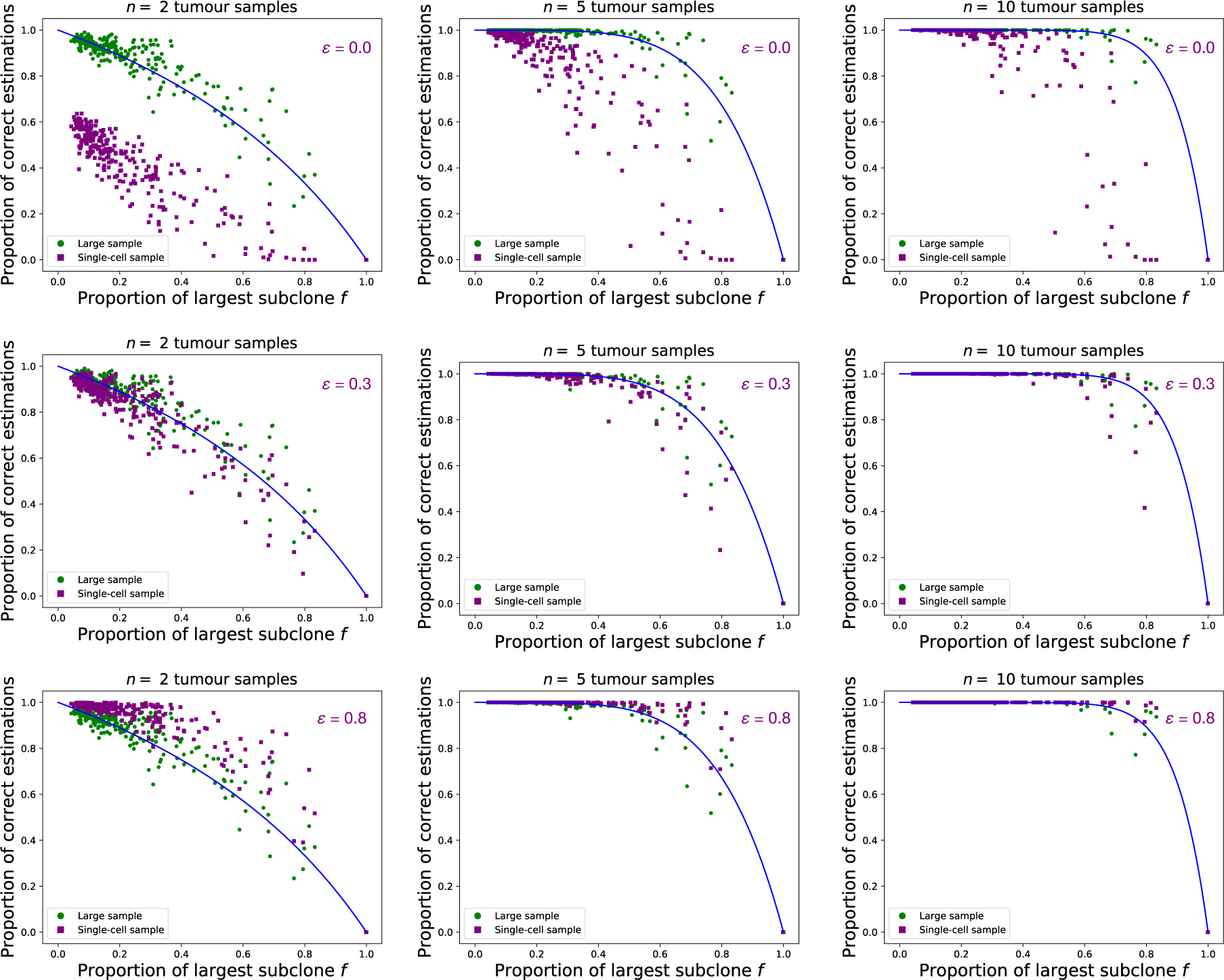
Clonality inference on the three-dimensional, spatial tumour model. To test the robustness of our results, we repeated the clonality inference process in the previously published spatial cancer model of Waclaw et al. [32], which is very different from our model. Small biopsies (green; one biopsy = 1 node) have a much greater probability to correctly classify clonal mutations than large biopsies (purple; one biopsy =8% of the total tumour size) if we include all mutations present in each sample *ε* = 0.0. As we increase the threshold of the mutation frequency *ε* (middle panel *ε* = 0.3, bottom panel *ε* = 0.8), the accuracy of larger samples is increasing, and goes beyond single-cell samples and model predictions, same as in our simpler model. model. Number of tumours = 300 with maximum size of 5 *·* 10^6^ cells were simulated with a death rate of *d* = 0.8. Mutation rate *γ* = 0.02 mutations per division.

**Figure 6:**
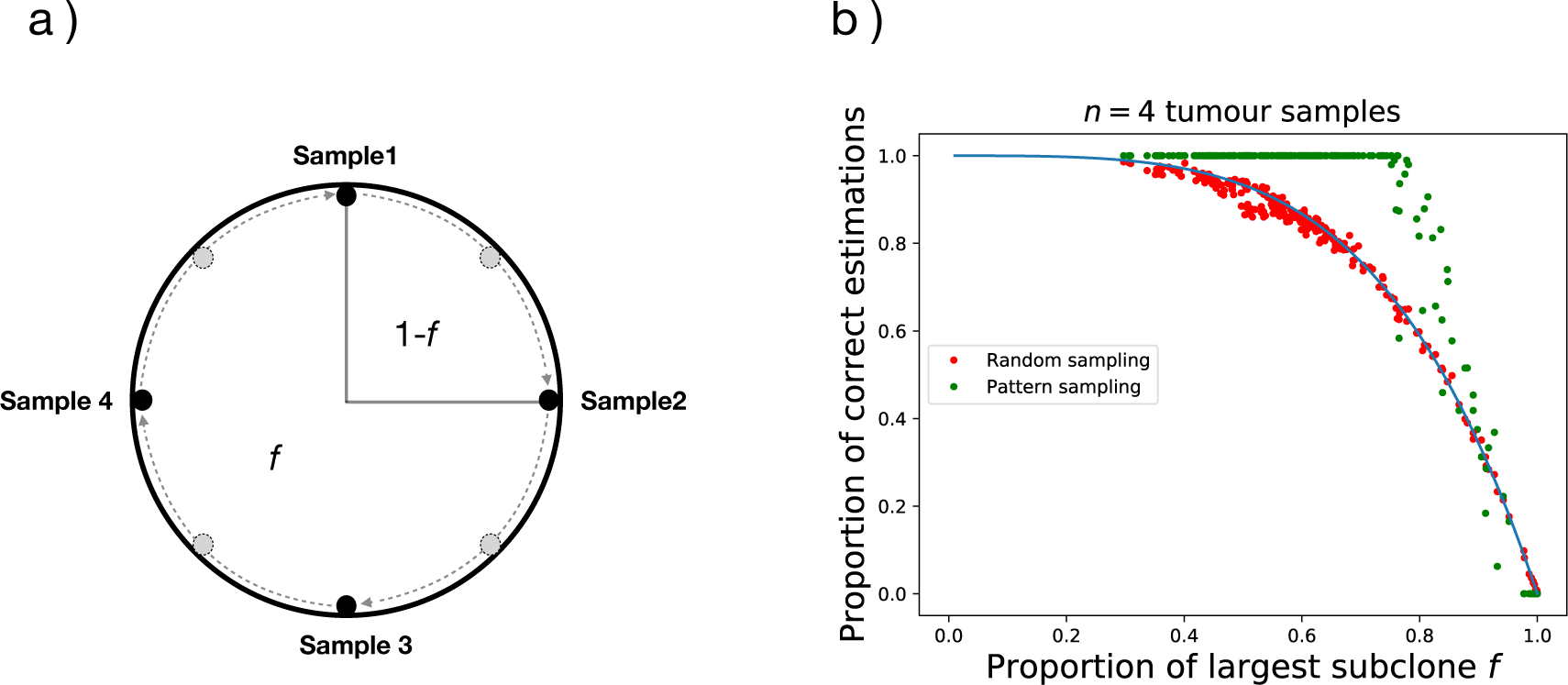
Pattern spatial sampling and identification of clonal mutations. We compared our model model (blue line) to simulation results using random single-cell sampling (red points) and to simulation results using sampling in specific pattern (blue points). Sampling pattern in panel a). Each red point represents the proportion of correct estimates with *n* = 4 samples after 10 000 sampling repetitions. For the green points, the template pattern was rotated to obtain multiple sampling repetitions from each tumour, while maintaining the distance between the cells and the peripheral location of cells. The proportion of correct estimations was calculated from a number of possible sampling repetitions for each tumour. Sampling in pattern appears superior to random sampling throughout most of the range of proportion of largest sub-clone *f* (Fig b)).

### Mathematical model

Let us first consider a simple model with only a single bifurcation representing the entire phylogenetic tree of the tumour. This bifurcation generates a branching subpopulations of cells that diverged directly from the ancestral population of initiating tumour cells. This branch contains a new sub-clonal mutation compared to the ancestral population. Here we define a balancing factor *f* as the proportion of the subpopulation within this branch, while the proportion of the other branch of the ancestral population is 1 − *f*. If we take *n* independent samples at random, the probability *pf* (*n*) of finding the true clonal (truncal) alteration is the probability that not all *n* samples come from the branch with the new sub-clonal mutation, in our case this is

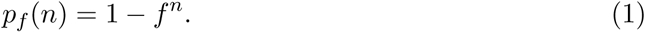

We now generalize the expression of *pf* (*n*) for a phylogenetic tree with a large number of bifurcations. Among all bifurcations, we are specially interested in the branches that diverged directly from the ancestral population. These branches are defined as first-tier branches, each of which contains a distinct sub-clonal mutation compared to the ancestral population (Figure 1d). Within each of these first-tier subpopulations, subsequent sub-clonal mutations would happen constantly (Figure 1d). However, these subsequent events are unnecessary for identifying the truly truncal alteration, so in what follows we ignore them and focus on the first-tier branches. We assume that the mutation rate of cells in the ancestral population does not change over time, so the first-tier branching mutations arrive after equidistant intervals. As a result, the balance factor *f* is unchanged and applies to all the first-tier branching mutations. Suppose there are *M* first-tier branches, which are ordered by their time of occurrence. The proportion of cells in the *k*th first-tier subpopulation is *f* times (1 − *f*)^*k*−1^ (the fraction of cells that did not carry any of the previous *k -* 1 first-tier sub-clonal mutations). In this way, the proportions of cells in these first-tier branches are given by *f*, (1 − *f*)*f*, (1 − *f*)^2^*f*, …, (1 − *f*)^*M*−1^*f*. To identify the true truncal alteration with *n* independent samples, these samples should not come from one single first-tier subpopulation. Thus, the probability *pf* (*n*) is given by

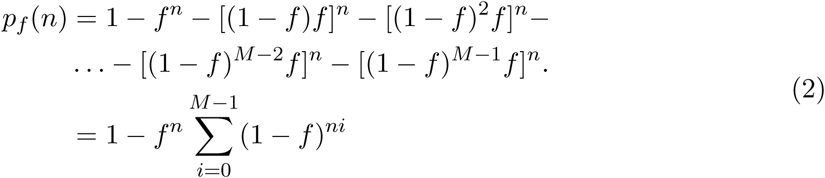

If *M* is sufficiently large, the geometric series can be used to approximate *pf* (*n*) by the simplified expression

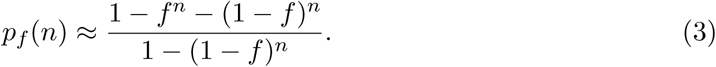

The parameter *f*, necessary for the calculation of the probability *pf* (*n*) can be estimated from data in the following way: The idea is to compare the sets of clonal mutations identified by all permutations of tumour samples. More specifically, first the intersection of all alterations of all *n* biopsy samples has to be determined. After that, the intersection of all possible combinations of *i* = 2 biopsy samples is generated. The frequency at which both intersections coincide is an estimate for the probability of a correct classification, *p*_*f*_ (*i*)/*p*_*f*_ (*n*). The same procedure is performed for all possible combinations of *i* = 3, 4, *…n* − 1 biopsy samples. Using these probabilities, one can estimate *f* for a given cancer by fitting the estimated probabilities to

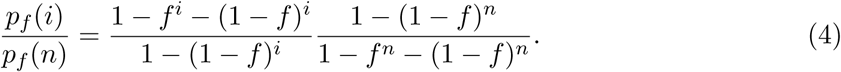

## Results

### Expectation from our mathematical model

In a highly homogeneous cancer with low mutational burden, even the largest sub-clonal mutation is only present in a small proportion of the tumor. Thus, our mathematical model predicts a fairly small chance to get false positive clonal mutations, see Fig. 2a with *f* = 0.1. Already with *n* = 3 samples the probability to correctly classify truly clonal mutations is *>* 98%. For the case where the first branching event leads to a tumour with two roughly equally-sized populations *f* = 0.5 (both of which will carry a tremendous amount of private mutations and many subsequent branchings) we reach a probability of *>* 98% already with *n* = 6 samples. Finally, in tumours where specific sub-clonal mutations undergo great expansion, it is highly likely that this expanding mutation and its sub-clonal mutations will be categorized as clonal (Fig. 2a with *f* = 0.9). This is because with increasing *f* it becomes less probable to sample from the part of the cancer without that abundant sub-clonal mutation. To reach the same level of confidence *>* 98%, as in shown tumours with lower *f*, we need *>* 38 samples for *f* = 0.9.

### Validation of the mathematical model using single cell sampling in simulated tumors

The proportion of the largest sub-clonal mutation *f* has a great effect on the clonality analysis. The probability to correctly classify clonal mutations with *n* = 2, *n* = 5 and *n* = 10 independent samples changes substantially with the value of *f* (Figure 2 b.) In principle, it is possible to correctly estimate clonality with only two samples, in particular if the largest sub-clone is sufficiently small. Theoretically, two samples give a correct estimation with probability *>* 98% for tumours where the proportion of the largest sub-clonal mutation is below *f* = 0.1. However, this probability drops rapidly with increasing *f*. Using more samples can substantially increase the probability of correct clonality assessments. With *n* = 10 samples, we cover most of the range of *f* and only for *f >* 0.8, our estimates become less reliable. For most values of *f*, 10 random tumour samples are sufficient to reach a high probability of a correct clonality assessment. We validate our mathematical model by comparing it with the results from stochastic spatial simulations of cancer growth (Figure 2b for *n* = 2, 5, 10.). Each point represents the proportion of correct estimations of clonality inferred from 10 000 iterations of *n* independent and random samples from a single tumour, in which the proportion of the largest sub-clonal mutation is *f*. Results obtained from simulations are in good agreement with our theoretical expectation, in particular results with more than two samples show almost perfect agreement between the mathematical model and the simulated tumours, despite the spatial correlations between clones arising in our computational model – which become crucial if we sample more than one cell.

### Effect of the sample size on clonality estimation

Biological samples from tumours typically contain a variety of different subpopulations of cancer cells, healthy surrounding and ‘supporting’ tissue as well as leukocytes that infiltrate the tumour. All of these cells can influence the interpretation of the sequencing data and the correct assignment of mutations. In current clinical applications, sequencing a group of cancer cells is standard – with single-cell genomic profiling so far an approach of the future. For that reason, we investigated the clonality inference by using multiple large samples, each containing 1% of the total number of cells in the tumor. In the previous analysis of single cell samples, there was no mutational frequency component – every mutation was of equal value for the clonality estimation. With multiple cells, we gain additional information of the frequency for each mutation within the sample.

#### Well-mixed tumours

As we previously stated, in our analysis a sub-clonal mutation is misclassified as clonal if it is present in all samples, therefore to classify it correctly there should be at least one sample where that mutation is absent. In a well-mixed tumour, mutational frequencies within large single samples already represent the spectrum of frequencies within the whole tumour. That makes them unusable for the classification of truly clonal mutations from multiple samples by means of exclusion from multiple sampling, as a large number of mutations appear clonal if we do not consider the frequency of each mutation within the sample. We can reduce the number of candidate mutations by excluding mutations with frequency below a certain threshold *ε*. By doing so, we remove sub-clonal mutations with low frequencies from the analysis and get a high proportion of correct estimates (Figure 3). Bringing the threshold *ε* above the frequency of the most abundant sub-clonal mutation (*f*) leads to a correct clonality assessment regardless of the number of samples. However, such clear demarcation likely is the result of our idealized scenario. In reality copy number changes and limited sequencing depth shift frequencies of mutations and introduce additional errors, leaving some uncertainty for the minimal list of clonal mutations.

#### Structured tumours

In structured tumours the clonality inference with multiple large samples is less accurate than the equivalent single-cell sample when including all mutations in the analysis, even though with large samples much more cells are included in the analysis than with single-cell sampling (Figure 4 for *ε* = 0). Large samples (1% of tumour size) contain more sub-clonal mutations that might be considered clonal. Having the frequency of each mutation within the sample, we can consider all mutations present at sufficiently low frequency within the sample as sub-clonal. By doing so we stop considering all mutations present in every sample as clonal. By raising the mutation detection threshold *ε*, low frequency sub-clonal mutations are removed from the analysis which increases the accuracy of our classification, ultimately surpassing the probabilities predicted by our model for single-cell sampling (Figure 4). We would get the best results by considering only mutations that appear clonal within the sample. Yet, some of clonal mutations might appear sub-clonal within the samples due to contamination with healthy tissue, copy number variation or sequencing noise, and would be wrongly excluded from the analysis.

To further test our approach, we repeated the same inference on a spatial model of tumour growth originally developed by Waclaw et al. [32], shown in Fig. 5. This model is very different from ours not only in the dimensionality, but also in many other details of the stochastic implementation. For example, the model is not based on a spatial lattice, allowing more complex configurations of cells in space. Nonetheless, the results on this three-dimensional model show the same qualitative features we observed in our two-dimensional scenario. The structure of the tumour has the same effect on the probability to correctly detect clones, furthermore both size of the samples and removal of sub-clonal mutations within samples from the analysis are showing similar trends as in our original computational implementation.

### Clonality inference using non-random spatial sampling

Until now, we only considered a random sampling process. In reality, this approach is not applicable and it would be useful to have clearly defined spatial relations between individual samples. Thus, we now compare the clonality inference using four samples arranged in a circular spatial pattern (Figure 6a) against four randomly distributed samples. We use samples from the (circular) edge of the tumour with the greatest distance between samples. In order to calculate the probability for a correct estimation of clonal mutations for a single tumour, we repeat the sampling process on the same tumour after we rotate all samples (Figure 6a) while maintaining the distance between them. We find that by sampling in this pattern, we increase the probability to correctly classify clonal mutations in tumours across most range of sub-clone proportions, *f*, compared to random sampling (Figure 6b). This is especially pronounced for intermediate number of samples. Interestingly, for *n >* 1*/*(1 − *f*), the classification of clonal mutations is correct almost with certainty. Only in cases where the proportion of the largest sub-clone *f* is close to 1, random sampling can be superior to pattern sampling.

These results can be generated to any number of samples positioned in an equidistant pattern. If we keep the same distance between samples, to have at least one sample not containing the largest sub-clonal mutation (1 − *f*), on average we need to sample *n* = 1*/*(1 − *f*) samples. This translates to at least 10 samples needed for a correct classification of clonal mutations for tumours with *f >* 0.9.

To test the possibility that improvement in classification is caused by specific (unknown) properties of cells from the edge of tumour and not due to pattern sampling, we took a series of random samples from the edge of the tumour and estimated clonality. These results match those from purely random sampling.

## Discussion

Targeting “driver mutations” in cancer is considered an important new approach to therapy that takes into consideration a varied mutational landscape present in tumours, even when arising in the same tissue [34–36]. The paradigm set by TKI therapy of CML and ALK inhibition in a small subset of patients with non-small cell lung cancer is quite compelling. In these two tumours, the mechanistic understanding of how the mutation drives the tumour is clear and therefore the term “driver” mutation is justified. Similarly, c-Kit expression on gastrointestinal stromal tumours (GIST) renders these tumours sensitive to imatinib as are the rare cases of mastocytosis with eosinophilia due to PDGFRA expression or mutant c-Kit expression [37–39]. BRAF^V600E^ mutations in malignant melanoma render the cells sensitive to vemurafenib [40]. Sequencing of other rare tumours has also led to the discovery of mutations that can be meaningfully targeted [41]. However, in the majority of cases, the identification of a mutation by itself does not imply that it is a driver – even if this was shown to be the case in a similar tumour and it is present in a significant fraction of the sample. Clonality needs to be proven with a reasonable certainty if there is any hope that targeted therapy will be effective. Sequencing a single sample and inferring that a mutation is “actionable” is fraught with problems, since the sample of the tumour sequenced may not be representative of the whole tumour, and in addition sampling has also to contend with the problems of false positive and negatives, high background noise due to the potential presence of not fully malignant cells that may still harbor normal copies of important genes such as TP53 [42] as well as contamination with normal tissue. It is therefore not surprising that despite major efforts, the practical benefit of NGS sequencing for the individual patient to date has been limited. A recent example illustrates this case: In a series of 95 patients with cancer seen at MD Anderson Cancer Center, NGS sequencing identified at least one mutation in 92 % of patients. The most common were in TP53 (25%) and KRAS (10%). In principle, 36% of the tumours sequenced had an actionable mutation and 13 patients received therapy based on this sequencing data to target the presumed driver mutation. Four patients had a partial response, six had stable disease while three progressed [36]. It is difficult to justify the current clinical approach with these results. Proving that a mutation is truncal and therefore clonal should lead to better identification of driver mutations and proper targeting of such mutations is more likely to give meaningful results. It appears that a proper sampling strategy for multi-region sequencing of a tumour is a key component in the process for the correct identification of truly clonal mutations. Such a list that is developed for every unique patient sequenced will likely be enriched for ‘driver’ mutations. In this work, we discuss how to improve the strategy determining this list of clonal mutations with a high level of certainty. The future introduction of multi-region tumour profiling into clinical practice requires a better understanding of the underlying mechanisms of intratumour heterogeneity and a more standardized approach to tumour sampling. We are still unable to steer the biological processes within a tumour to affect its heterogeneity [43], but we can optimize the way we collect and analyze tumour samples.

Our study provides insights into both intrinsic and extrinsic factors that influence the probability to detect truly clonal mutations. As the construction of the complete tumour phylogeny is not necessary for clonality inference, we focused on the reconstruction of the series of branching events coming from an ancestral population. We have developed a mathematical model for the calculation of the probability for correct identification of truly clonal mutations from multi-region sampling of cancer with a large number of bifurcations. In that process, the largest sub-clone is the most relevant factor in the identification of truly clonal mutations. Its proportion is a consequence of the time since the emergence of the first sub-clone. The earlier the first sub-clonal mutation occurs, the more likely we are to misclassify it as truncal. A large abundance of this first sub-clonal mutation requires more samples to ensure that at least one sample does NOT contain that mutation. In addition, if the first sub-clone is only present in a small proportion of the tumour, there is a low probability of it being misclassified as clonal. Our results show that considering multiple branching events we now see that the probability to correctly classify mutations is much greater than previously thought [20].

In solid neoplasms, where cells grow in space, we have shown that larger samples are more likely to overestimate clonality of some mutations if the analysis include all mutations present in each sample. It is necessary to exclude mutations present in low and medium frequency from the analysis. Doing so we not just reach the same accuracy as we would by single-cell sampling, but we even increase it substantially. However, exclusion of lower frequency mutations from the analysis might cause false negative classification of some clonal mutations whose frequencies are variable within a sample due to contamination with healthy tissue, duplication of genetic material within some cancer cells, or sequencing error. In the presence of other detected clonal mutations, a false negative error is of less concern than a false positive, as false positives would deprive the patient of effective therapy. Decision on the cutoff for the mutation exclusion must be individually chosen based on the number of available clonal mutation candidates.

Finally, we bring a theoretical rationale for sampling in non-random patterns. We showed that placing biopsies in pattern equally distant from each other, might substantially increase the probability to correctly classify truly clonal mutations when compared with random sampling. We are aware that our computational simulations of tumour growth are simplified and lack many features of living systems, such as cell migration in tumours with more complex growth pattern. There, a different spatial sampling strategy might be more successful. Our results provide a rationale for pathologists when taking samples for multi region tumour sequencing and clinicians during endoscopic sampling of neoplasms of e.g. gastrointestinal tract. By choosing samples with maximum spread in suggested pattern one maximizes the chances to correctly classify clonal mutations. We also offer a way to estimate a level of certainty for a list of detected clonal mutations which can serve as a guidance to oncologists in their choice of appropriate target. Our results provide some considerations for the improved clinical assessment of targetable mutations in treatment naive tumours.

## Conclusions

In conclusion, the correct classification of clonal (truncal) mutations is of great importance for the success of anti-tumour therapy. We have shown how the probability to identify truly clonal mutations depends on sub-clonal composition of tumour and how many samples one must take to be able to discern mutation clonality with great confidence. Furthermore using a computational model of cancer heterogeneity, we have shown how the size of biopsies affects the probability to correctly identify clonal mutations. Finally, we showed that our suggested spatial sampling pattern is superior to random sampling.

## Abbreviations

TKI: Tyrosine kinase inhibitors
GIST: Gastrointestinal stromal tumour
CML: Chronic myeloid leukemia
ALK: Anaplastic lymphoma kinase
NGS: Next generation sequencing

## Declarations

## Availability of data and materials

Code available from corresponding author on request.

## Author’s contributions

All authors developed the concept, L.O. implemented simulations and analyzed the model with input from all authors. L.O. and A.T. wrote the paper with input from D.D., B.W and D.Z.. All authors read and approved the final manuscript.

## Acknowledgments

We thank Bartlomiej Waclaw for inspiring discussions and explanations on his model and Jordi Arranz and Marvin BÖttcher for programming support.

